# When large-effect damaging alleles enter small populations

**DOI:** 10.1101/2025.09.11.675554

**Authors:** J. Ingemar Ohlsson, Sofia Mikko, Susanne Eriksson, Anna M. Johansson, Martin Johnsson

## Abstract

Large-effect recessive damaging variants found at relatively high frequencies are common enough in domestic animals that they cause problems in breeding programs. There are several potential reasons. Deleterious variants may rise in frequency because of genetic drift, be driven to intermediate frequency by balancing selection by heterozygote advantage, or be spread by exceptionally influential parents, usually sires, that sometimes occur in animal breeding programs. Further, animal populations are often connected by migration. In this paper, we use a series of population genetic models to address how large-effect damaging alleles establish themselves in small populations, such as domestic animal populations or wild populations of mammals. The results confirm that a lethal allele under directional selection is fairly unlikely to become common, unless there are influential sire events with very large contributions or the population size is very small. On the other hand, a lethal allele under balancing selection can easily become common, especially if there are influential parents. When there is balancing selection, migration between connected populations effectively spreads alleles. The results accord with the many examples of recessive damaging alleles in domestic animals.

## Introduction

In this paper, we use a series of population genetic models to address how large-effect damaging alleles establish themselves in small populations, such as domestic animal populations or wild populations of mammals. Such large-effect recessive deleterious variants at relatively high frequencies are common enough in domestic animals that they pose applied problems for breeders, and several population genetic processes can contribute to their establishment in a population.

In many cases, these damaging variants show evidence of balancing selection by heterozygote advantage (see reviews by [1, 2]. Wide-spread pleiotropy in combination with long-range linkage disequilibrium [3] and strong selection make heterozygote advantage an intuitively appealing explanation for the occurrence of damaging variants in animal breeding programs. Unfavourable genetic relationships have often been observed (reviewed by [4]) between production and fertility and health in farm animals or between desired morphological features and health in companion animals, suggesting that there is scope for causative variants to have antagonistic effects on multiple traits, which could manifest as heterozygote advantage.

However, there are also cases where a large-effect damaging variant is common but where no beneficial side-effect has been found [5, 6]. Furthermore, demonstrating heterozygote advantage at a single locus is not necessarily straightforward when there is population structure and potential confounding.

However, domestic animal populations are small enough that chance events may have large impact. In particular, influential parents, usually males, can drive deleterious variants to high frequency [7]. There are several known cases where particular deleterious alleles have been traced back to an influential sire, e.g., [8–10]. Such cases can be purely a matter of drift, if what influences the choice of sire is independent of the genotype at the locus in question, or a matter of linked selection due to beneficial variants elsewhere in the sire’s genome.

Further, migration between populations, including gene flow between breeds, may contribute to the spread of deleterious variants. Gene flow between populations has been historically important in breeds of domestic animals [11, 12], and currently has a large impact in breeds such as international cattle and horse breeds [13, 14].

There are several theoretical models that describe the behaviour of single-locus and two-locus systems affected by drift, selection and migration that can shed light on the potential contributions of these mechanisms. In this paper, we use population genetic models, including classical equation-based models and individual-based simulations, with the aim to analyse how a new damaging allele becomes common in a small population. We study the effect of heterozygote advantage, influential sires, and spread through migration.

## Methods

The scenarios modelled in this study include:

- selection on one or two loci, where the second has a beneficial effect,
- with or without heterozygote advantage for the damaging allele,
- with or without an initial influential sire effect,
- with or without migration exchange with a connected population,
- as idealised population simulations or stochastic simulations of individuals.

### Selection on one locus

The starting point for the analysis is the classical model of selection on a single locus, extended to balancing due to heterozygote advantage. In the classical model (see e.g., [15] or [16]), each genotype has a constant fitness value of *W*_*AA*_, *W*_*Aa*_ or *W*_*aa*_. The model proceeds in two steps over discrete generations. First, zygotes are formed through random mating, and then individuals reproduce according to the relative fitness of their genotype. To find the change in allele frequency in one generation of selection, we first calculate the relative fitness of each genotype weighted by their genotype frequency, assuming Hardy—Weinberg frequencies among the zygotes. Then, we sum the contribution of the two genotypes that include the allele of interest. This procedure gives a recurrence equation that can be iterated to calculate allele frequency over the generations. That is, in general:

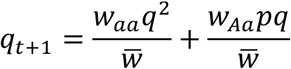

Where *p* is the current allele frequency of allele *A*, and *q* the current allele frequency of allele *a*, *q*_*t*+1_ is the frequency of allele *a* in the next generation, and *W*% = *W*_*AA*_*p*^2^ + 2*W*_*Aa*_*pq* + *W*_*aa*_*q*^2^ is the average fitness. We use this equation to calculate expected allele frequency trajectories from deterministic models. Going forward, we will take allele *a* to be the damaging allele. In cases without balancing selection, there will be directional selection against *a*, and in cases with balancing selection by heterozygote advantage, we assume that *a* also has some beneficial side-effect which gives the heterozygote genotype the highest net fitness.

#### Classical model without heterozygote advantage

For a damaging allele without heterozygote advantage, the fitness of the wildtype homozygote *AA* is set to *W*_*AA*_ = 1, and the fitness of the damaging allele homozygote can be parameterised as *W*_*aa*_ = 1 − *s*, and the fitness of the heterozygote as *W*_*Aa*_ = 1 − ℎ*s*, where *s* is the selection coefficient and *h* the degree of dominance. For simplicity, this paper will use fully recessive lethal alleles as an example in the main text, that is *s* = 1 and ℎ = 0. Results for other parameter combinations are presented in supplementary materials.

#### Classical model with heterozygote advantage

For a damaging allele under balancing selection due to heterozygote advantage, following Hedrick [1], the fitness of the heterozygote *Aa* is set to *W*_*Aa*_ = 1, and the fitness of the two homozygotes are expressed as *W*_*AA*_ = 1 − *s*_1_ for *AA* and *W*_*aa*_ = 1 − *s*_2_ for *aa*. (Hedrick called the selection coefficient *s*_2_′, because *s*_2_ was already used in another parametrisation; here, the prime is dropped.) For the purpose of this paper, “balancing selection” means balancing selection due to such heterozygote advantage, not for instance because of frequency-dependent fitness values or fluctuating selection.

Again, this paper will use alleles that have a fully recessive lethal effect as an example in the main text, that is *s*_2_ = 1. For a such an allele, the equilibrium frequency [1] is:

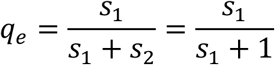

Most of the time we will assume that the equilibrium frequency *q*_*e*_ = 0.10, which corresponds to a selection coefficient *s*_1_ ≈ 0.11. This equilibrium frequency is similar to the frequency of several damaging alleles assumed to be under balancing selection [1], but one should keep in mind that allele frequencies vary between populations and that these alleles may not actually be at equilibrium. Results for other parameter combinations is presented in supplementary materials, and the R package provided with the paper allows analysing any parameter values.

#### Influential sire effect

For the simplest possible approach to model the introduction of a damaging allele into a population by an influential sire, imagine that the damaging allele occurs only in one influential individual who is a carrier of the deleterious allele and sires a certain proportion of *p*_*sire*_ of the offspring in the first generation. If we start counting from his offspring, this is setting the starting frequency to *q*_0_ = *p*_*sire*_/4, because the influential sire contributes one half of the chromosomes of the proportion he sires, and on average half of those offspring receive the deleterious allele. In the absence of beneficial alleles in the sire, i.e., when his large contribution is purely a matter of chance, the fate of the damaging allele is described by the classical models, only starting from a high *q*_0_ due to the influential sire effect. This model assumes that going forward from the influential sire event, there is random mating, and inbreeding is ignored.

#### Classical model with migration

To model migration between populations, we imagine two connected populations with symmetric migration rate *m*. We call the first population, where the damaging allele originates the “origin” population and the second population, that is free from the damaging allele at the starting time, the “recipient” population. We start from the classical model of a damaging allele, either with or without balancing selection by heterozygote advantage and add a migration step where surviving individuals move between populations, with consequences for allele frequency:

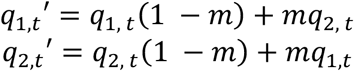

After having modified the allele frequencies by migration, we apply the same recurrence equations as for the classical models to *q*_1,_′ and *q*_2,*t*_′, within each population. These are the recurrence equations, with an additional subscript *i* = 1,2 to indicate the two populations and a prime to indicate that this is frequencies after it has been modified by migration. We allow for different genotypic fitness values in the two populations.

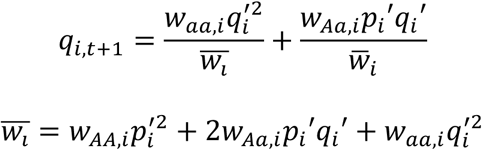

#### Classical two-locus model

When an influential sire became influential because of some beneficial allele that he is carrying, there can be linked selection between the damaging allele and the beneficial allele. Because the damaging and beneficial alleles co-occur within his genome, there will be linkage disequilibrium between them. This can be described by a two-locus model [17, 18] that tracks the frequencies *x*_1_, …, *x*_4_ of the four gametes *AB, Ab, aB,* and *ab*. The two loci are assumed to have recombination rate *r* between them, and we also track the coefficient of allelic association *r* = *x*_1_*x*_4_ − *x*_2_*x*_3_ which quantifies linkage disequilibrium. The recurrence equations for gamete frequency are:

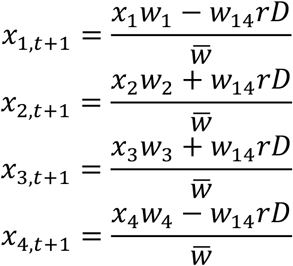

Where *W*_*i*j_ are fitness values for each genotype, *W*_*i*_ = ∑_j_ *W*_*i*j_*x*_j_ is the marginal fitness of gamete *i*, i.e., the frequency-weighted average fitness of all genotypes that contain the gamete, and 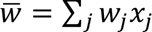 is the average fitness. *W*_14_, in particular, is the fitness of double heterozygotes.

We assume that the lethal allele is introduced into the population by a popular sire who also carries a linked, additive, beneficial allele of large effect. The size of the beneficial effect was set to be [inlin],, giving the double wildtype homozygote the same fitness as in the above heterozygote advantage models. We assume that the influential sire is homozygous for the beneficial allele, and that the beneficial allele has a frequency of *q*_0*B*_ = 0.1 among the other gametes. That is, the beneficial allele is common, but not yet carried by the majority of the animals, leaving scope for selection. A supplementary table includes cases when the beneficial allele is absent from the rest of the population.

Fitness values for the nine two-locus genotypes are:

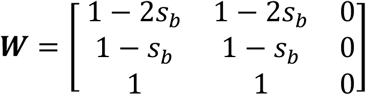

Where columns correspond to genotypes at the lethal locus (from left to right: *AA*, *Aa* and *aa*), and rows correspond to genotypes at the beneficial locus (from top to bottom: *BB*, *Bb* and *bb)*.

We calculate the frequencies from this model for with 2%, 5% or 10% offspring from the influential sire, ranging from large to very large contributions, and with recombination rates of 0.1%, 1%, 10% and 50%, ranging from closely linked to unlinked.

### Probability matrix models

We model the probability of a new mutation establishing itself in a population with a probability matrix model [15]. Probability (or transition) matrix models can be used to derive fixation probabilities and other parameters for Wright—Fisher models with genetic drift.

The idea is to set up a matrix of transition probabilities between all possible allele counts for a single locus in a population of a particular size. Each row corresponds to a state with a particular number of alleles *X*_*i*_ in the next generation and each column to the current number of alleles *X*_j_the current generation. The probability of going to from one state to is given by a binomial model with *2N* trials and success probability *Ψ*_*i*_.

Then, the probability of a particular state at a particular generation ***Y_t_*** can be calculated by matrix multiplication:

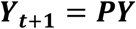

Where ***P*** is the probability transition matrix and ***Y*** is the current distribution of states. When modelling a new mutation entering the population, the initial distribution is taken to be one copy of the new damaging allele with probability 1 and probability 0 for all other states. We calculate the results for populations of size 100, 300, and 1000 over 100 generations.

Following Krukov et al. [19], we incorporate selection by making the binomial sampling parameter a function of fitness. In general, the parameter will be the frequency of allele *a* after selection, namely the contribution of each genotype weighted by relative fitness and standardized by average fitness:

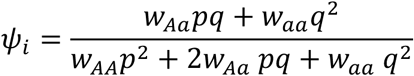

Specifically, this becomes 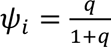 for a lethal allele without heterozygote advantage and 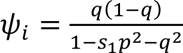 for a lethal allele with heterozygote advantage.

Alternatively, we use a probability matrix model of [20], designed specifically for lethal alleles. Their model allows for heterozygote advantage for a recessive lethal, and results in a probability matrix model similar to the above Wright—Fisher model, but with sampling from a Binomial distribution with *N* trials and success probability 2*Ψ*_*i*_, where the binomial sampling parameter is as above. Their original model also includes mutation, which we ignore here.

When modelling an allele being introduced by a popular sire, the initial distribution is the probabilities of the allele counts after random mating with a popular sire. These are derived from a binomial distribution with size *2N* and success probability equal to the contribution of the sire divided by four (*p*_*sire*_/4).

### Ideal population simulations

We also perform simulation of a population with a single damaging allele with and without heterozygote advantage. The model has discrete generations and a constant population size of *N* zygotes. The simulation happens in three steps for each generation:

1. Combine gametes from the previous generation into zygotes. For the starting population, the *2N* gametes are drawn from a Bernoulli trial with a success probability equal to the allele frequency, *X* ∼ *Binom*(1, *q*).
2. Perform selection of parents according to genotypic fitness values as in the classical models, drawing parents randomly with replacement with probability weighted by their relative fitness.
3. Sample one of the gametes each from the selected parents for each time they reproduce. This creates the gamete pool for the next generation. Calculate allele frequency among the gametes and record it.

To simulate a new mutation entering a population, we set the starting gamete pool to contain one copy of the allele, giving 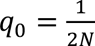 . We run these simulations with and without heterozygote advantage, for population sizes of 100, 300 and 1000. We replicate these simulations 5000 times.

#### Ideal population simulations with influential sire

We perform simulations of an influential sire effect in an ideal population using the same population-based simulation methods as above, setting the starting allele frequency based on the contribution of the influential sire *q*_0_ = *p*_*sire*_/4. We run these simulations with and without heterozygote advantage, for a population size of 300. We replicate these simulations 5000 times.

#### Ideal population simulations with migration

We also perform simulations of two connected populations using the same population-based simulation methods as above. We create two populations of equal size, each with sampling of zygotes and selection within population as above. However, between sampling and selection, we exchange *Nm* zygotes between populations to simulate migration with migration rate *m*. To model the introduction of a new mutation, we start with a gamete pool where the mutant allele exists only in one copy in one of the populations. As above, we run these simulations with and without heterozygote advantage, for population sizes of 100, 300 and 1000, and with migration rates of 0.01, 0.05, and 0.1. We replicate these simulations 5000 times.

### Two-locus simulations

We perform two-locus simulations to supplement the deterministic analysis of the linked selection case. In the two-locus model, we follow the frequency of the four gametes generated by two loci with two alleles each, and the fitness of the four possible two-locus genotypes.

The simulation happens in three steps for each generation analogous to the single-locus ideal population simulation above:

1. Combine two-locus gametes from the previous generation into zygotes. For the starting population, the gamete pool consists of one part from the influential sire and one part from other sires. For the part contributed other sires, there are 2*N*(1 − *p*_*sire*_) gametes that all carry the wild type allele at the lethal locus *A*, and randomly sampled alleles at the beneficial locus drawn form a Bernoulli trial, *X*_*B*_ ∼ *Binom*(1, *q*_0*B*_). Out of the 2*Np*_*sire*_ gametes contributed by the influential sire, half carry the mutant lethal allele *a* and half the wild-type allele *A* in combination with the favoured allele *b* at the beneficial locus.
2. Perform selection of parents according to genotypic fitness, using the same fitness matrix as in the two-locus model above, and drawing parents randomly with replacement based on probability weighted by their relative fitness.
3. Sample one of the two-locus gametes each from the selected parents for each time they reproduce. With probability given by the recombination rate *r*, the alleles at one locus will be switched between the haplotypes to produce a recombinant gamete. This creates the gamete pool for the next generation. Calculate allele frequencies for both loci and the linkage disequilibrium coefficient *D* = *p*_*ab*_ − *q*_*A*_*q*_*B*_ between the mutant alleles and record them.

We run these simulations with and without heterozygote advantage, for population size of 300 with 2%, 5% or 10% offspring from the influential sire, and with recombination rates of 0.1%, 1%, 10% and 50%. We replicate these simulations 5000 times.

### Breeding simulations

We also perform breeding simulations of an influential sire event in a more complex population structure using AlphaSimR [21]. We use a breeding structure inspired by the Swedish Warmblood sport horse population, with 300 sires and 3000 dams each generation, generating 6000 foals, half of each sex. The distribution of offspring of the sires was based on numbers of offspring from Swedish Warmblood sires 2008–2017. We have used this model before; see [22].

For the purpose of this paper, we make three changes to the model. Previously, we modelled the ongoing split within the warmblood sport horse population between horses selected for show jumping and those selected for dressage by simultaneous selection for two different breeding goals. Here, we simplify the model by selecting only on one quantitative trait.

Previously, we also used a distribution of different genetic effects for the allele under balancing selection, drawing them from a normal distribution. Here, we set the additive genetic coefficient of the selected variant at an effect size that approximately corresponds to *s*_1_ ≈ 0.11. Previously, we assumed that the lethal allele started out at an allele frequency around 0.05, similar to the frequency observed in Swedish Warmblood horses. Here, to model the introduction of a new mutation, we introduce the mutation in one copy in the most influential male in generation 5, and follow the population for another 40 generations.

We set this additive genetic coefficient with a method analogous to how additive genetic coefficient can be approximately converted into selection coefficients under truncation selection [16]. For a fully lethal allele, there are only two genotypes in the adult population, wildtype homozygotes and heterozygote carriers. The heterozygote is taken as the reference genotype with a fitness value of one. By definition, the average genetic difference between genotypes is equal to the additive genetic coefficient *a*. For a given selected proportion of heterozygotes *p*, the difference in proportion selected between the genotypes is approximated by a rectangle of area *aip*, where *i* is the selection intensity, assuming that the phenotypes are normally distributed and that the difference between genotypes is small compared to the standard deviation. This leads to an expression for the selection coefficient 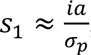, which can be rearranged to:

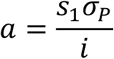

Where the selection coefficient *s*_1_ = 0.11, the phenotypic standard deviation *σ_P_* is set at the simulated value and *i* = 0.877, which is the sex-averaged selection intensity when 10% of the males and all females are selected.

In this model, we assigned the distribution of offspring by dividing sires into deciles, and assigning the most offspring to the top decile and increasingly fewer to the following, with equal numbers of offspring for each sire within a decile. This was based on the real distribution of offspring in Swedish Warmblood horses. The sires in the highest decile each had 116 offspring, meaning that the most influential sire makes a contribution of 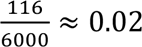 To simulate an influential sire event in the first generation, we simulate cases where the most influential sire has 5% percent and 10% the offspring, adjusting the rest of the offspring distribution proportionally. For the following generations, we used the unmodified distribution, where the most influential sire only gets 116 offspring as above.

The effective population size for this simulated population is around 300 based on the equation for effective population size with variable offspring numbers [15]:

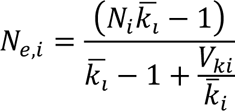

Where 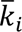 is the mean offspring number, *V_kix_* is the variance of offspring number, and *N*_*i*_ is the number of parents, and the index *i* indicates males or females. The sex-specific effective population sizes are then combined with 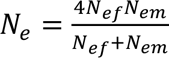.

We simulated two selection scenarios: one where the damaging allele was genuinely pleiotropic leading to heterozygote advantage and where the damaging allele has no effect on the trait, and there is only declining linkage disequilibrium with the beneficial alleles carried by the influential sire. We replicate these simulations 100 times.

#### Breeding simulations with migration

We extend our breeding simulation to migration in a similar fashion as the classical model, by adding a second population that has the same breeding structure and genetic background, and is connected to the first by a mutual migration rate. Before selection, a number of males, defined by migration rate *m* times the population size, are exchanged between populations.

We assume that females remain in their population and that only males are exchanged, similar both to domestic animal breeding programs and male-dispersal animals in the wild. We vary the migration rate, expressed in fraction of sires exchanged, with values 0.02, 0.1 and 0.2.

These are twice the values used in the ideal population simulation, because they are limited to males.

We introduce the damaging variant in one of the populations by an influential sire event where the allele arises in one copy in the most fit sire in the first generation, and trace the effect on allele frequency in both populations. As above, the influential sire event either occurs in the most fit sire for a total of around 2% of the offspring (116 offspring), or we increase his influence to 5% or 10% for the generation following the mutation event.

As above, we simulated two selection scenarios: one with genuine heterozygote advantage on the allele itself and one without, where beneficial alleles carried by the influential sire are merely in linkage disequilibrium with the damaging allele. We replicate these simulations 100 times.

### Software implementation

The models and simulations used in the paper consist of an R package that implements the equation-based models and ideal population simulations (https://github.com/mrtnj/selectionr) and a set of scripts that run the particular analyses used in the paper (https://github.com/mrtnj/balancing_selection_models). The two-locus models use Rcpp [23] to interface with C++ code to speed up simulation. The breeding simulations use AlphaSimR [21] version 1.3.4.

## Results

### The probability that a damaging variant becomes common with and without heterozygote advantage

Probability matrix models and ideal population simulations both showed that the probability that a new lethal mutation establishes itself in a population was low, unless there was balancing selection by heterozygote advantage. Figure 1 shows ideal population simulations of a lethal allele entering a population, and the probability of observing a lethal allele at a frequency above 0.05 from simulation and probability matrix calculations. On the other hand, with balancing selection, recessive lethal alleles were fairly likely to be observed at frequencies above 0.05. While the probability of observing the lethal allele at this frequency declined over time when the ideal population size was 100, probability still remained around 10% over tens of generations. In larger populations, establishment became more likely. There was a substantial lag between the introduction of the mutation and when it had increased to a frequency enough to meet the 0.05 threshold. In larger populations, that lag was longer.

**Figure 1.**
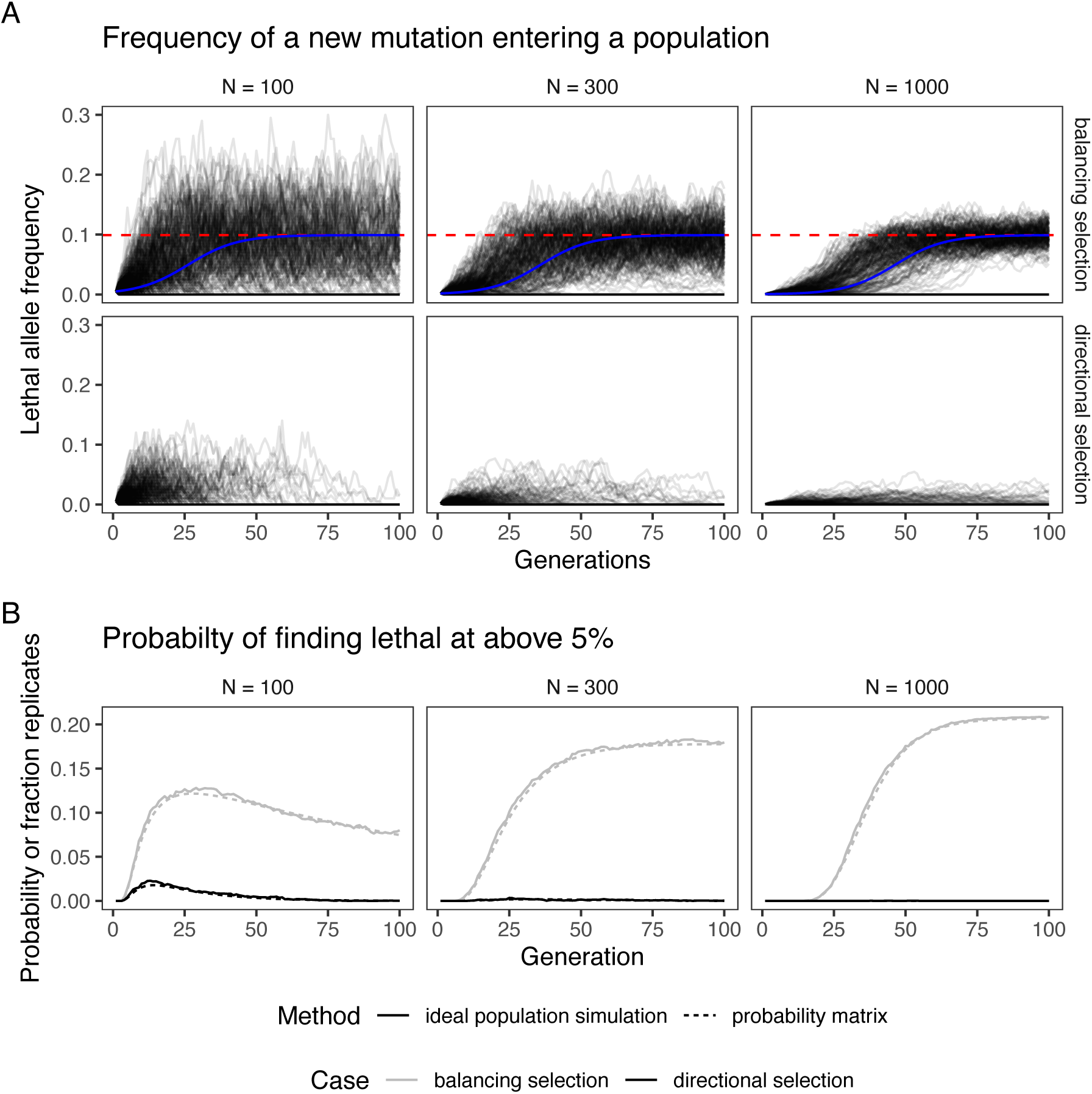
A new lethal allele entering a population with and without balancing selection by heterozygote advantage. A) Ideal population simulations of a new recessive lethal mutation entering a population. The lines show allele frequency trajectories of simulation replicates. The panels show the cases with population sizes of 100, 300, and 1000 individuals with and without heterozygote advantage for the lethal allele. The blue curve shows the allele frequency trajectory of the classical deterministic model, and the dashed red line shows the equilibrium frequency. B) Probability of a new recessive lethal mutation being observed at high frequency in an ideal population. The dashed lines show the probability of the lethal allele being at a frequency higher than 0.05 in a given generation, calculated from a probability matrix model. The solid lines show the fraction of simulation replicates where the lethal allele is at a frequency higher than 0.05 in ideal population simulations.

Probability matrix models based on the Wright—Fisher model and the model of [20] gave similar results (Supplementary Figure 1). Supplementary Figures 2 and 3 shows a wider range of parameters, varying the selection coefficients, using the probability matrix model. These illustrate the large influence of the strength of positive selection in the case of balancing selection, and the large effect of dominance coefficient for a lethal allele under directional selection.

### Influential sire effects and linked selection

After an influential sire event, the probability of observing a lethal allele at high frequency declined over time when there was directional selection but showed a plateau when there was balancing selection. Figure 2 shows the probability of observing a lethal allele at a frequency above 0.05 after an influential sire event in the case of a population size of 300. With balancing selection and 5% contribution, the probability of establishment went above 50%.

**Figure 2.**
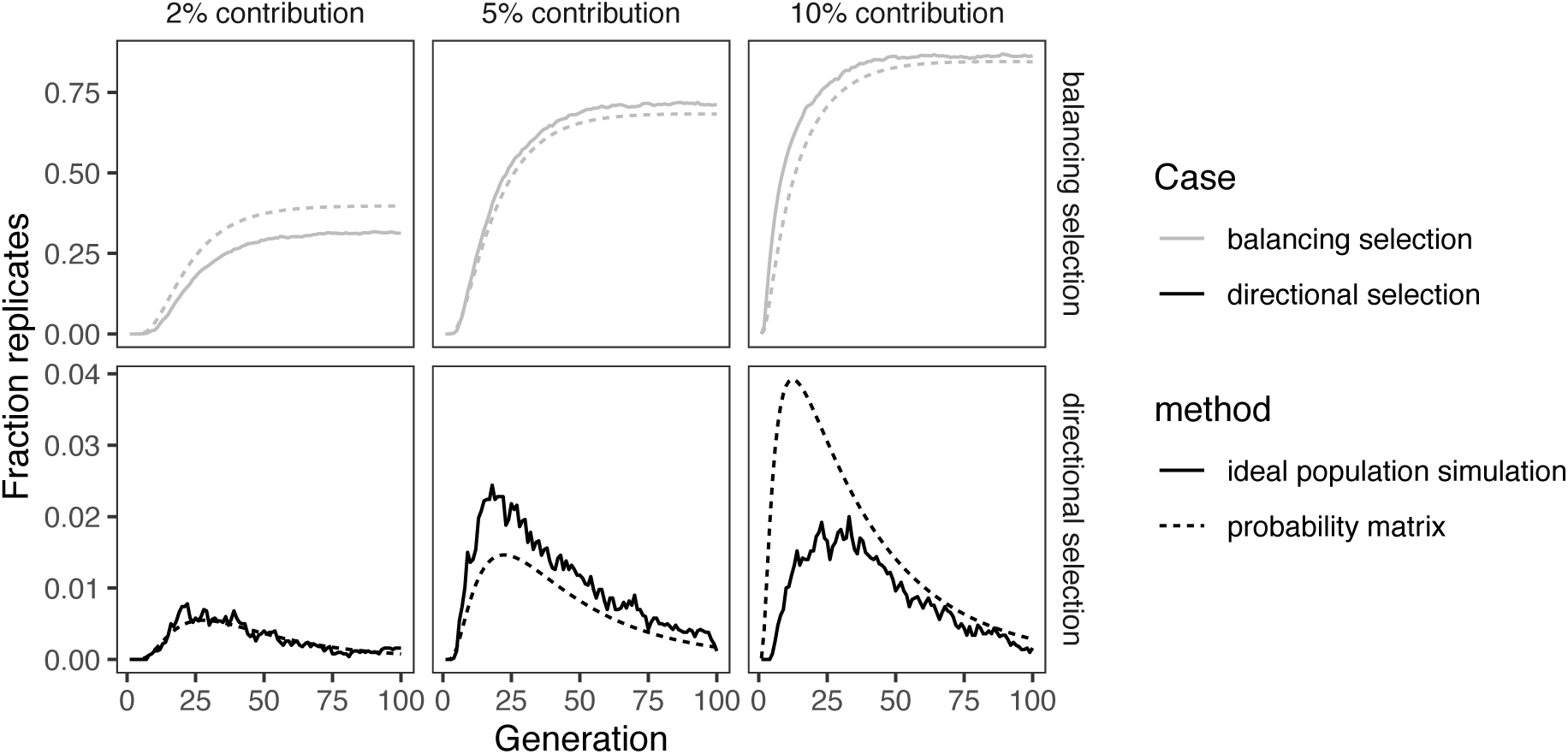
Probability of a new recessive lethal allele being observed at high frequency in an ideal population when introduced by an influential sire with population size N = 300. The dashed lines show the probability of the lethal allele being at a frequency higher than 0.05 in a given generation, calculated from a probability matrix model. The solid lines show the fraction of simulation replicates where the lethal allele is at a frequency higher than 0.05 in ideal population simulations. The panels show the cases with 2%, 5%, and 10% contributions of the influential sire.

However, even with 10% contribution from the influential sire, the probability of observing a lethal allele without balancing selection at frequency above 0.05 remained less than 4%.

Supplementary Figures 3 and 4 shows a wider range of parameters, varying the selection coefficient against the damaging allele and the population size using the probability matrix model.

A more complex breeding simulation (Figure 3), based on the Swedish Warmblood sport horse population, showed similar patterns. A lethal allele without balancing selection was unlikely to be observed at high frequency, whereas a lethal allele with balancing selection was likely to establish itself. In the case of 10% contribution from the sire, establishment of the allele was virtually certain. However, as predicted by classical models and shown in the breeding simulation, if a lethal allele introduced by a popular sire by chance reaches a high frequency, it takes a long time for its frequency to decline.

**Figure 3.**
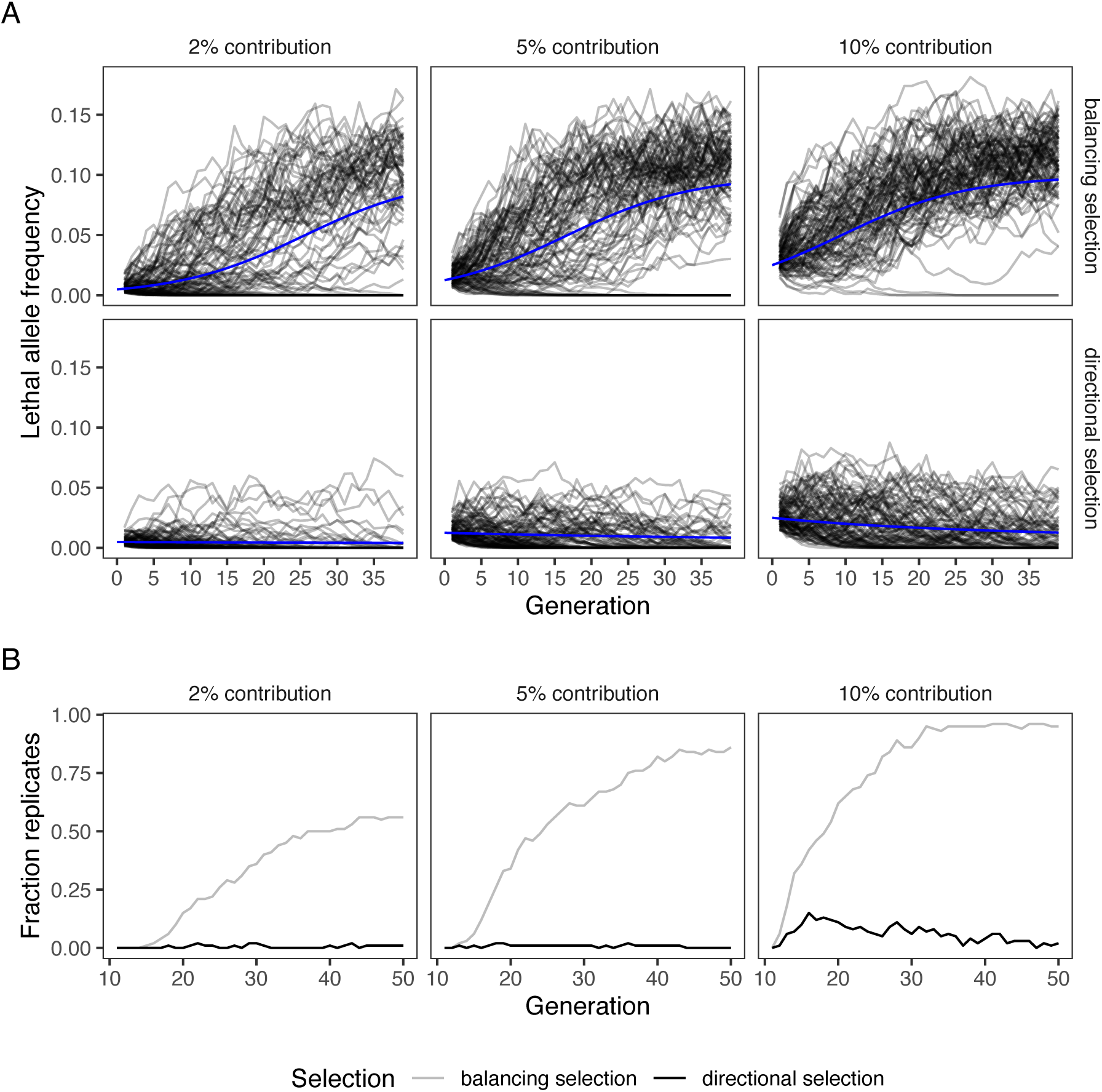
Breeding simulation of a new recessive lethal mutation introduced by an influential sire. A) Frequency trajectories of simulations with and without balancing selection. The blue lines show trajectories from the classical deterministic model for with selection coefficient corresponding to the simulated additive genetic coefficient. B) Probability of a new recessive lethal mutation introduced by an influential sire being observed at high frequency with and without balancing selection. Lines show the fraction of simulation replicates where the lethal allele is at a frequency higher than 0.05 in a breeding simulation with an influential sire effect.

Beneficial linkage disequilibrium from introduction by a selected influential sire was able to maintain a damaging variant at high frequency for a long period. Figure 4 shows the change in allele frequency due to a linked beneficial variant and the probability of observing the lethal allele above a frequency of 0.05 after an influential sire event for different recombination rates in the case of a population size of 300. The probability of observing the lethal allele at frequency above 0.05 was qualitatively similar to the case of a small population with balancing selection, where the probability decreased over time but remained substantial for tens of generations. For example, with 5% contribution and low recombination rate between beneficial and lethal variants, the probability of observing the lethal at above 0.05 peaked at around 10%.

**Figure 4.**
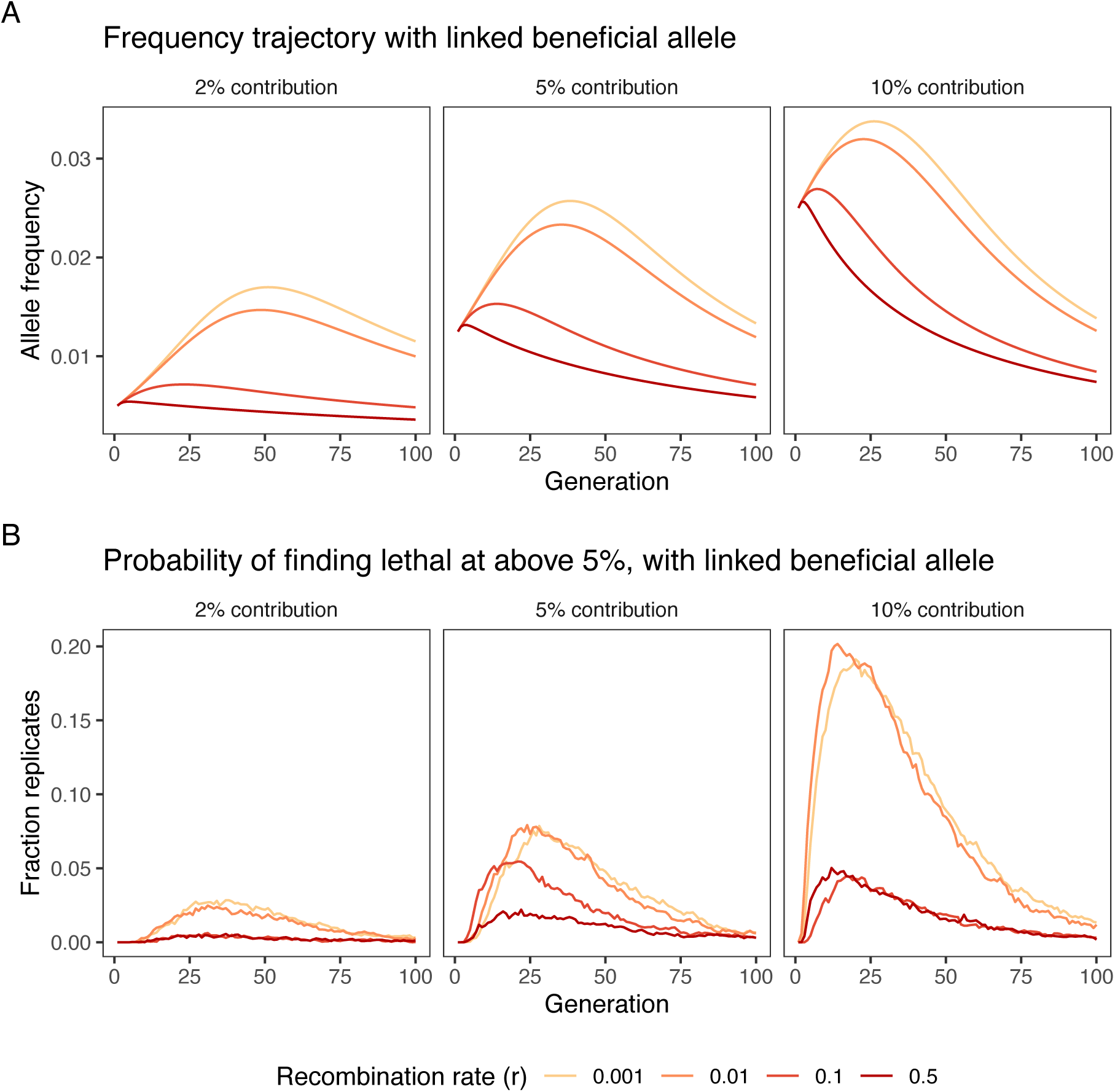
A) Allele frequency trajectories from deterministic two-locus model. B) Probability of a new recessive lethal mutation being observed at high frequency when linked with a beneficial allele. Lines show the fraction of simulation replicates where the lethal allele is at a frequency higher than 0.05 in a two-locus simulation. The colours represent recombination rate. The panels show the cases with 2%, 5%, and 10% contributions of the influential sire.

Supplementary Table 1 shows the predicted number of generations for the frequency of a lethal allele to decrease from 0.025 to 0.005 after an influential sire event with a linked beneficial variant, compared to a recessive lethal without linked beneficial variants. When recombination rate was high, the expected time was almost the same, but when recombination rate was low and the beneficial allele absent from the rest of the population, the decrease took about 100 generations longer.

### Spread by migration between connected populations

Migration between connected populations was able to effectively spread a damaging allele with heterozygote advantage. Figure 5 shows the probability of observing a recessive lethal mutation in two connected populations, derived from ideal population simulations. Figure 6 shows the same for an increase due to an influential sire event. Compared to the single-population case, migration promoted the establishment of a lethal allele with balancing selection in the originator population, and at the same time delayed it. When migration rate was high, the allele established itself more slowly than it would with low migration, but the probability of establishment was in the end similar or higher. In the recipient population, the establishment happened more slowly, but when the population migration rate was high, it was about as likely as in the originator population.

**Figure 5.**
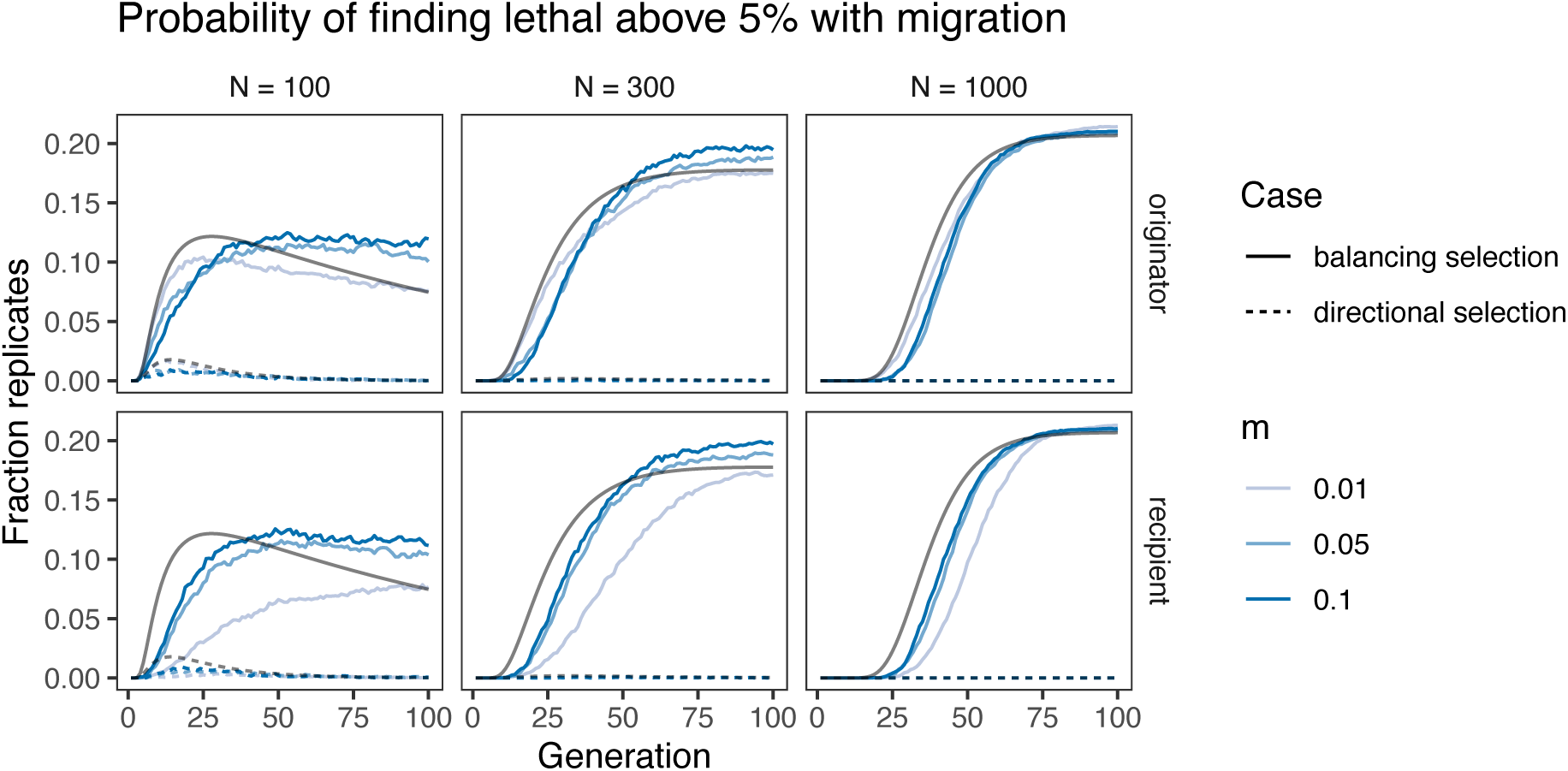
Probability of a new recessive lethal mutation being observed at high frequency in two populations connected by migration. The lines show the fraction of replicates where the lethal allele is at frequency higher than 0.05 in ideal population simulations of two connected populations. Dashed lines are lethal alleles without balancing selection; solid lines are lethal alleles with balancing selection. The colours represent different migration rates. Panels show frequency in the originator and recipient populations, and cases with population sizes of 100, 300, and 1000 individuals per population. The superimposed grey lines show the single-locus predictions from a probability matrix model (i.e., the same curves as Figure 1).

**Figure 6.**
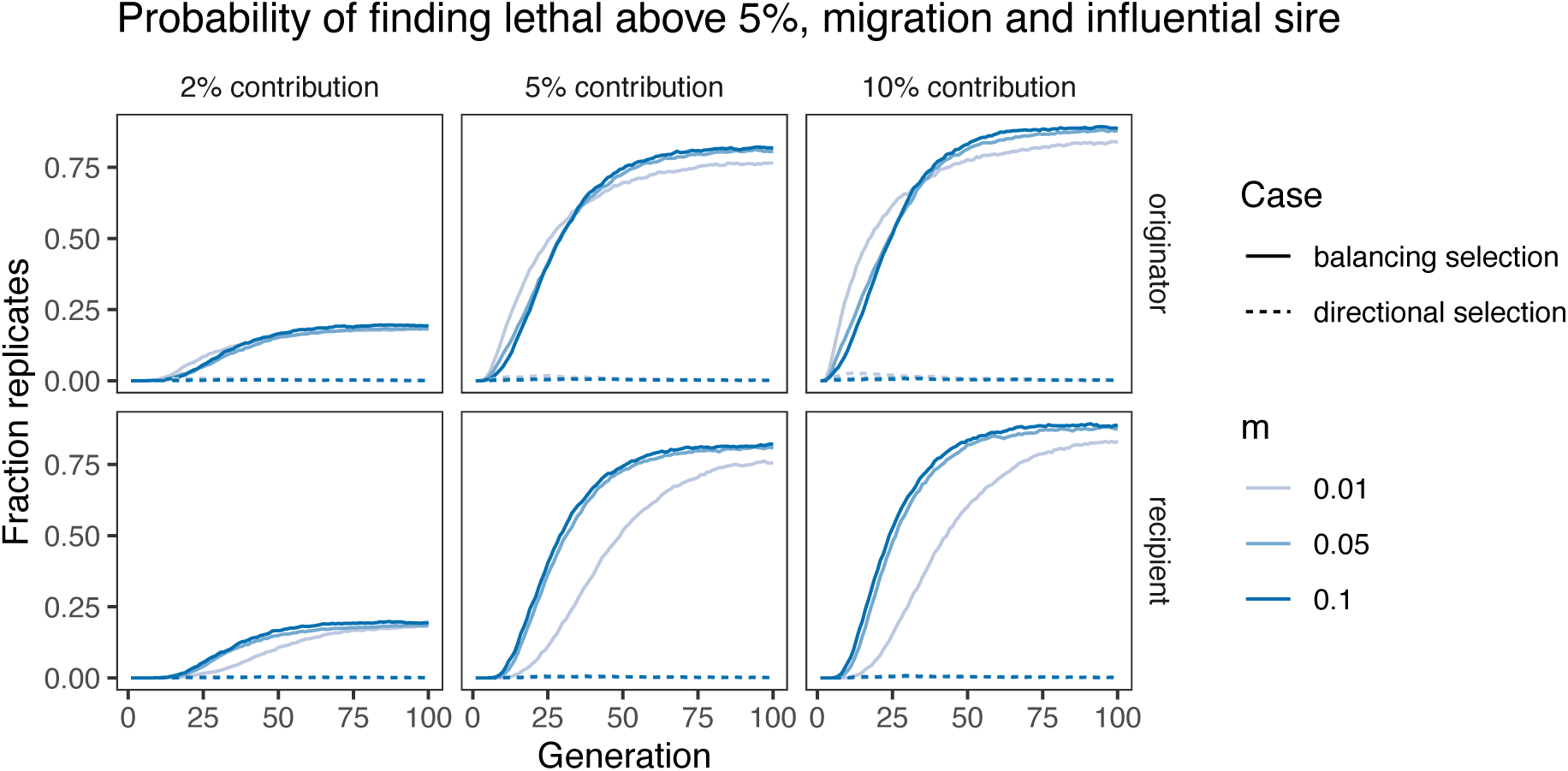
Probability of a recessive lethal allele being observed at high frequency in two populations connected by migration after an influential sire event. The population size of each population was 300. The lines show the fraction of replicates where the lethal allele is at frequency higher than 0.05 in ideal population simulations of two connected populations. Dashed lines are lethal alleles without balancing selection; solid lines are lethal alleles with balancing selection. The colours represent different migration rates. Panels show frequency in the originator and recipient populations, and cases with 2%, 5%, and 10% contributions of the influential sire.

### Probability of finding lethal above 5% with migration

The trajectories in allele frequency when the damaging allele was introduced in connected populations differed qualitatively from that of an isolated population. Supplementary Figure 6 shows the predicted allele frequency trajectories from the classical deterministic model for a lethal allele with and without heterozygote advantage in two connected populations. In the recipient population, the allele frequency after an influential sire event first increased by migration, before decreasing again.

In the case without heterozygote advantage, migration to and from a population without the lethal allele helped decrease allele frequency. Supplemental Table 1 shows the predicted number of generations for the frequency of a lethal allele to decrease from 0.025 to 0.005 with and without migration. The decrease happened 40-50 generations faster.

Heterozygote advantage in the originator population could also drive establishment of the lethal allele in a recipient population where there was no positive selection. Supplementary Figure 7 shows the same ideal population simulation as Figure 6, except with no positive selection in the recipient population, meaning that the allele acts like a recessive lethal in this population.

Again, more complex breeding simulation (Figure 7), based on the Swedish Warmblood sport horse population, showed a similar result, where a lethal allele with heterozygote advantage effectively spread between connected populations due to migration, in this case only of sires. The increase in probability of observing the allele in the recipient population was delayed by up to 20 generations compared to the originator population.

**Figure 7.**
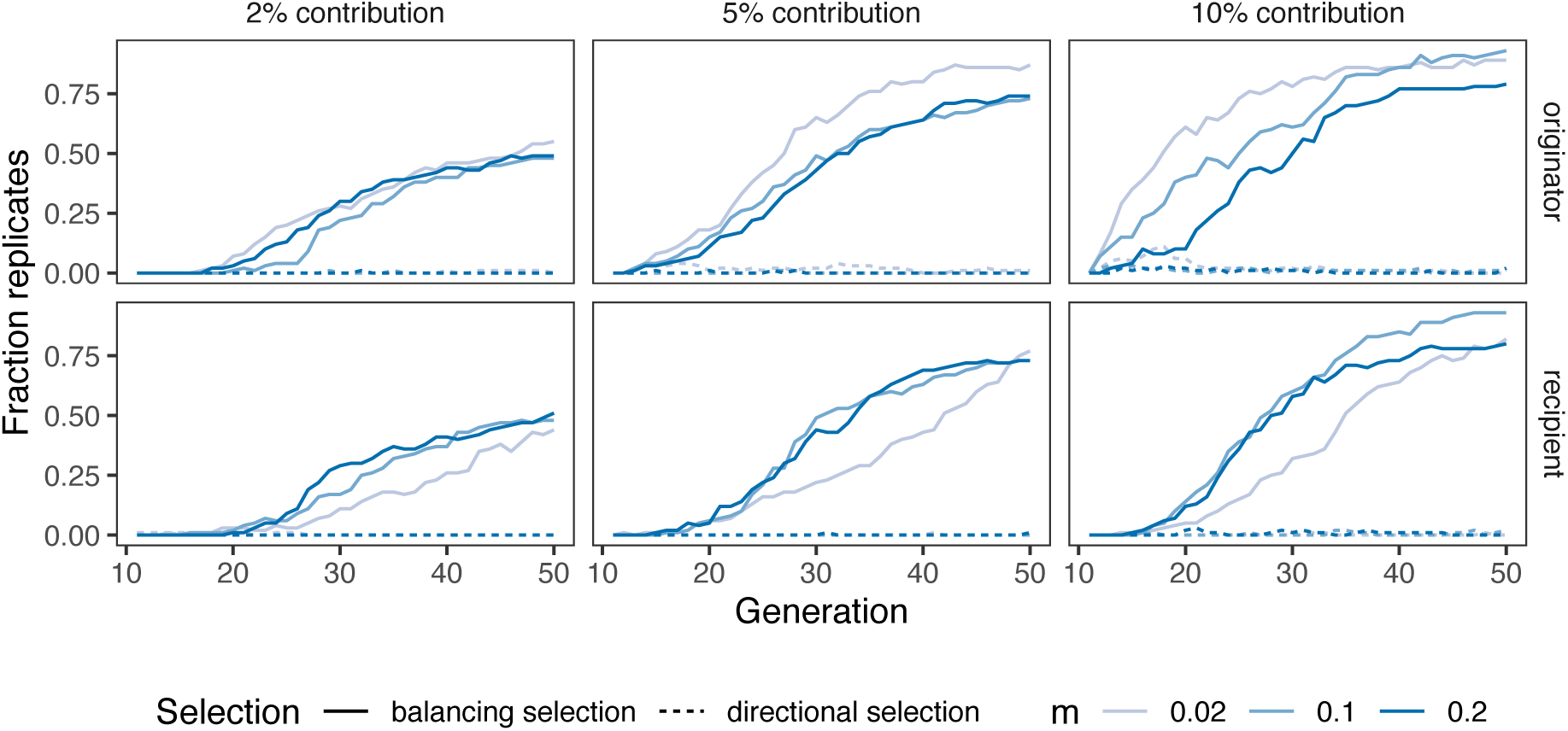
Breeding simulation of a new recessive lethal mutation introduced by an influential sire, with migration. The lines show the fraction of replicates where the lethal allele is at frequency higher than 0.05. The colours represent different migration rates. Panels show frequency in the originator and recipient populations, and cases with 2%, 5%, and 10% contributions of the influential sire.

## Discussion

This paper analysed the behaviour of allele frequencies of large-effect deleterious variants in small populations, such as farm and companion animals, with and without heterozygote advantage, influential sire events and migration. We address the overarching question of what processes drive large-effect deleterious variants to high frequencies in small populations. The results confirm that a lethal allele without balancing selection is fairly unlikely to become common, unless there are influential sire events with very large contributions or the population size is very small. A lethal allele with heterozygote advantage, on the other hand, can easily become common, especially if there are influential sire events. Lethal alleles with heterozygote advantage were also effectively spread between populations by migration.

Based on these results, we will now discuss the prevalence of heterozygote advantage in small populations, the transient effect of influential parents, the effect of migration, and finally compare our results to real cases in horses, cattle and pigs.

### The prevalence of balancing selection by heterozygote advantage

These models confirm that that even in small populations, balancing selection due to heterozygote advantage makes a lethal allele much more likely to establish itself than it would be due to drift alone, in agreement with the many known or suspected cases of damaging variants under balancing selection in domestic animals (see examples below). However, the resulting prevalence of balancing selection among common damaging variants also depends on the relative mutation rates of balanced and non-balanced damaging mutations, which is difficult to know.

Partial penetrance, resulting in large-effect but sub-lethal damaging alleles also establish more easily than recessive lethal alleles. Estimates of the effect distribution of new mutations from humans and model organisms suggest that most deleterious variants have small effects, [24, 25] and also that partial dominance is common [26]. Theoretical models as well as empirical examples suggest that pleiotropic effects are also common, which gives scope for balancing selection [27].

Further, in effectively small populations, like domestic animals, linkage disequilibrium extends over relatively long range [3]. This means that transient heterozygote advantage due to linkage is likely. In our models, there can be a substantial effect of linkage between beneficial and damaging variants when the recombination rate between them was 1%. This association will break up over time, but with the generation time of many domestic animals, tens of generations could be tens to hundreds of years.

### The transient effect of an influential parent

Once a lethal, or sub-lethal, allele has risen in frequency through drift or an influential sire event, it decreases in frequency only slowly as can be seen from the classical model and our simulations of an influential sire event. In both ideal population and breeding simulations, there are replicates where a lethal allele without heterozygote advantage rises to appreciable frequency through drift, and stays there for a long time. Even without balancing selection, the loss of damaging variants is slow and subject to large fluctuations, compared to the time scales that we are observing in a domestic animal population. The models suggest that influential sire events are able to make large-effect deleterious variants common, and that they are expected to remain common for a long time, especially when there is beneficial linkage.

This is like a form of transient heterozygote advantage.

Because of the limited time scales that are relevant to domestic animals, one can question the value of equilibrium-based models for understanding deleterious variants in farm and companion animals. We do not dispute the logic or validity of equilibrium-based arguments, but note that the questions that farm and companion animal geneticists and breeders need to ask often have to do with the fate of particular alleles currently segregating in populations, not about the long-term equilibrium behaviour.

Therefore, one important limitation of this work is the assumption of constant population size after the occurrence of the mutation or after the influential sire event. For many animals with systematic breeding programs, there have been recent, dramatic, changes in effective population size, so that few populations are likely to have been at near constant size over a hundred generations [28, 29]. Our models could be extended to population histories that change over time.

### Migration efficiently spreads damaging variants with balancing selection

In practical terms, our results present a case for international collaboration around monogenic traits in animal breeding programs. In many species, animal breeding is international with gene flow between national populations and, to some extent, between breeds. Both cattle and horses have strong international exchange of genetics. While migration impeded the spread of a recessive lethal without balancing selection, it promoted the spread of damaging variants with heterozygote advantage.

Migration may spread a damaging allele even if there is not positive selection in the recipient population. Our previous work on the fragile foal syndrome allele in the two diverging specialties within the Swedish Warmblood sport horse can be seen as an example. Previously strong but decreasing connectedness between show jumping and non-show jumping horses [30] and positive selection in non-show jumping horses appears to have driven a lower, yet still substantial, frequency of the lethal allele in the show jumping population, where there is little evidence of positive selection on the allele [22].

After a popular sire event, migration also complicates the picture by introducing transient counter-intuitive increases and declines in allele frequency. This occurs when the influx of carrier animals via migration into the recipient population overcomes the change due to selection, and the influx of mutation-free animals overcomes selection in the originator population. These patterns are evident in predicted allele frequency trajectories from classical deterministic models, but, in a small population, these patterns are liable to drowning in the noise of genetic drift.

### Real cases in horses, cattle and pigs

Using the fragile foal syndrome allele [31], which inspired the breeding simulations in this study, as an example, this allele is widely spread between warmblood and related horse populations and therefore likely to be old. The presence of the fragile foal syndrome allele in multiple populations [32], connected by gene flow, fits better with the expected behaviour under balancing selection than a lethal without balancing selection. There is evidence of balancing selection in the form of differences in performance-related traits between carriers and non-carriers [22]. However, the generation interval is around 10 years [33], meaning that any transient effect due to an influential sire event, linked selection or migration will take a long time to resolve.

In cattle, there are several examples of recessive deleterious alleles that have an association with milk production traits indicating that balancing selection have kept these alleles in the population despite the negative effects for homozygotes (Cole *et al.*, 2016; Georges *et al.*, 2019; Jenko *et al.*, 2019). A very clear example of balancing selection in cattle is a 660 kb deletion that leads to embryonic lethality in homozygous state but that also has a positive effect on milk yield in heterozygotes and have quite high frequencies in Nordic red cattle breeds [38]. A mutation causing muscle weakness in calves is an example of where a recessive mutation can be traced back to a sire used in breeding several decades ago [37].

In pigs, many lethal recessive alleles were identified by [39]. Examples of balancing selection involving direct meat production trait in pigs is given by [40] where a mutation causing leg weakness in homozygotes is associated with increased muscle depth in heterozygotes, and by [41] where a mutation causing lethality in homozygotes are associated with increased growth rate in heterozygotes. Another study shows an example of how a mutation that are lethal in homozygotes can still be maintained in the population since heterozygotes have an improved mothering ability compared to pigs that do not have the mutation at all [6]. Mothering ability is important in pigs since a large litter size at birth is insufficient if piglet mortality until weaning is high.

These examples from horse, cattle and pig demonstrates that there are several identified examples of balancing selection in domestic species. There will most likely be many more examples in the future.

## Conclusions

In this paper, we use a series of population genetic models to analyse how a new damaging allele becomes common in a small population. Influential sires can spread large-effect damaging alleles effectively and give rise to a transient balancing selection effect due to beneficial alleles carried by the sire. Migration between connected populations effectively spreads damaging alleles with heterozygote advantage. These results illustrate how large-effect damaging alleles can become common in populations, and how short-term allele frequency changes can be poor guides to the underlying selection. In the process, we have implemented the classical models and ideal population simulation in an R package that may be used for parameter exploration and teaching.

## Acknowledgements

This work was supported by Formas – a Swedish research council for sustainable development (Dnr. 2020-01637) and by the Kjell and Märta Beijer Foundation. We thank Åsa Gelinder Viklund for fruitful discussions.

---

**Supplementary Figure 1.**
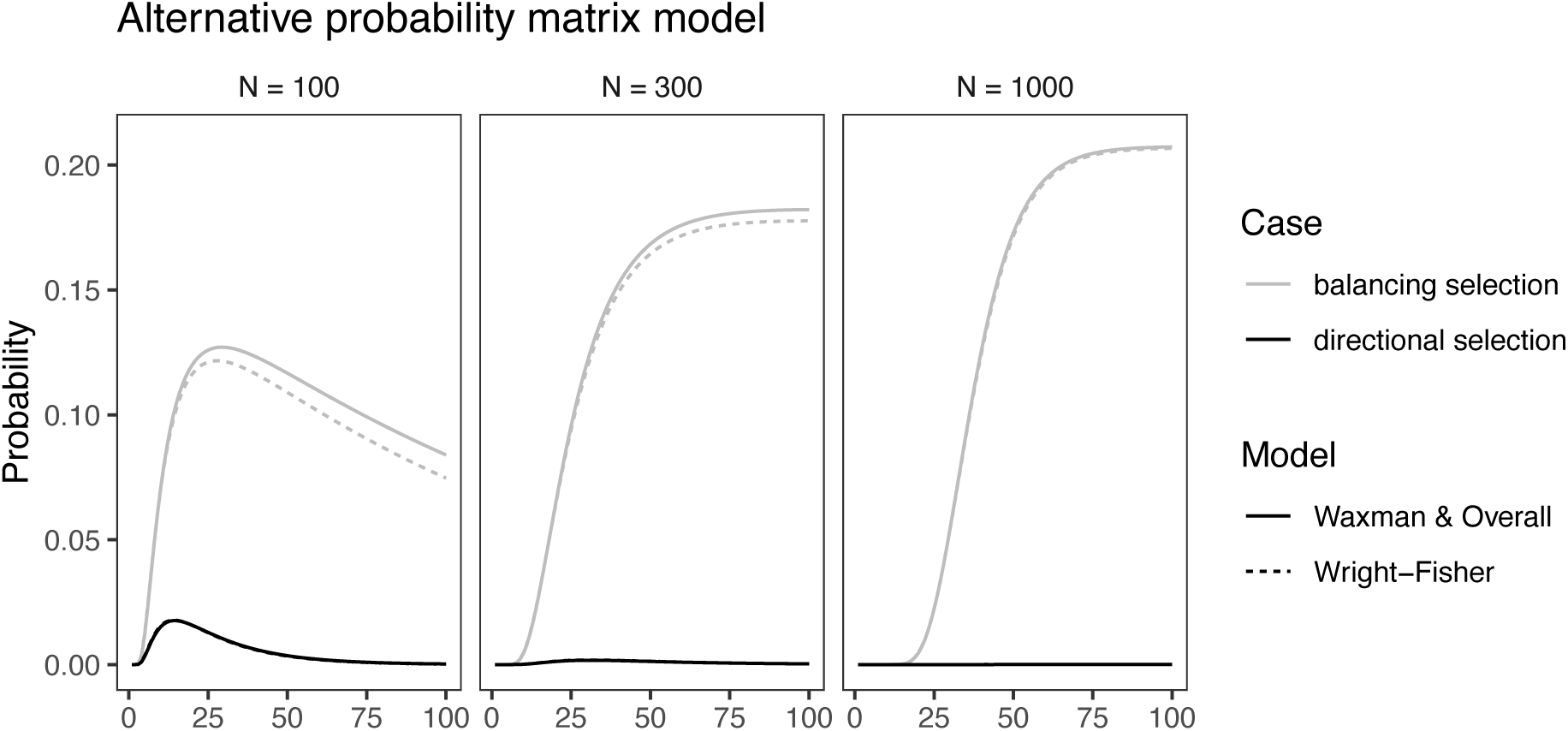
Comparison of probability matrix calculations using a Wright— Fisher population and the model for lethal alleles by Waxman & Overall.

**Supplementary Figure 2.**
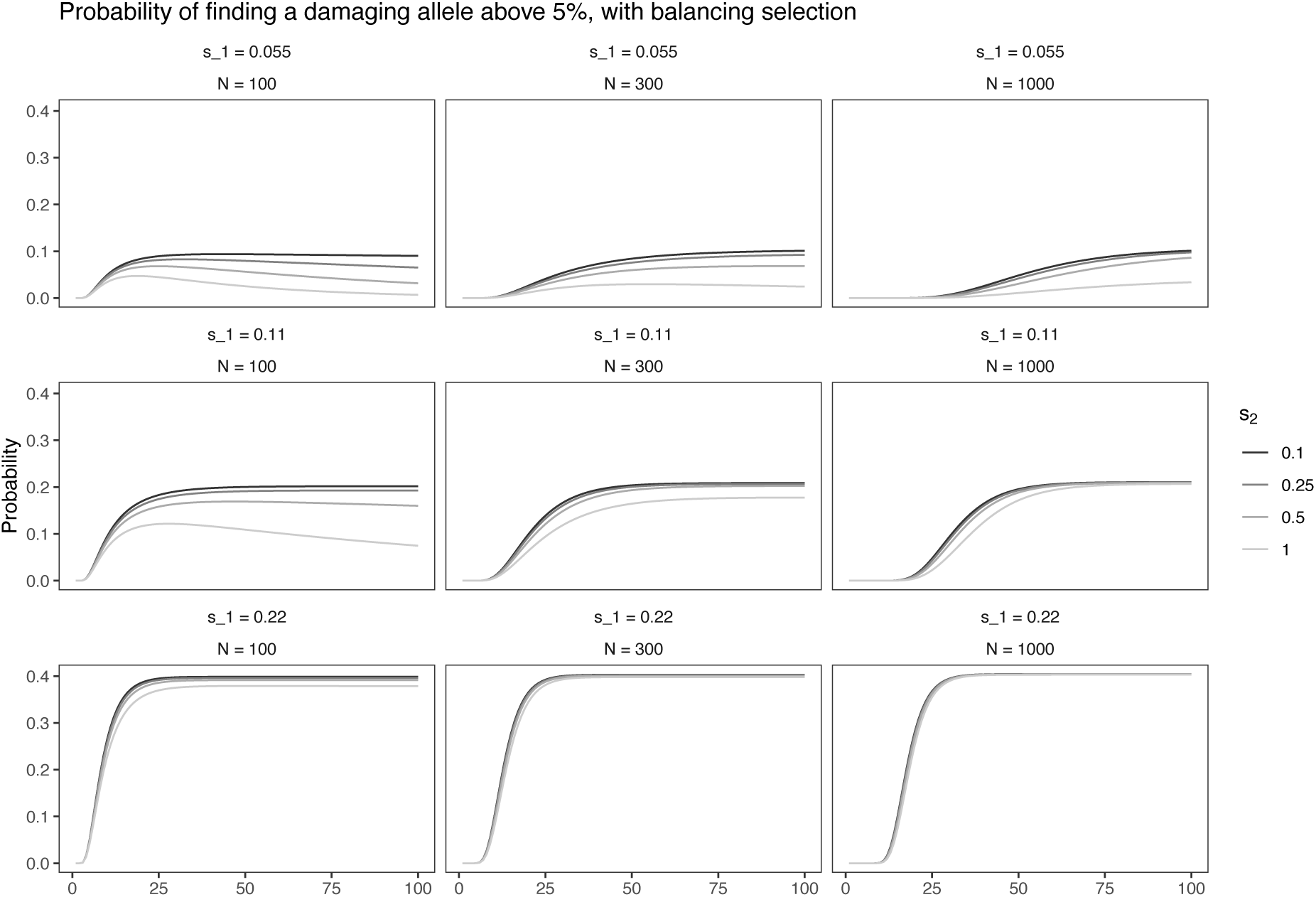
Parameter exploration for damaging alleles with heterozygote advantage, using the probability matrix model.

**Supplementary Figure 3.**
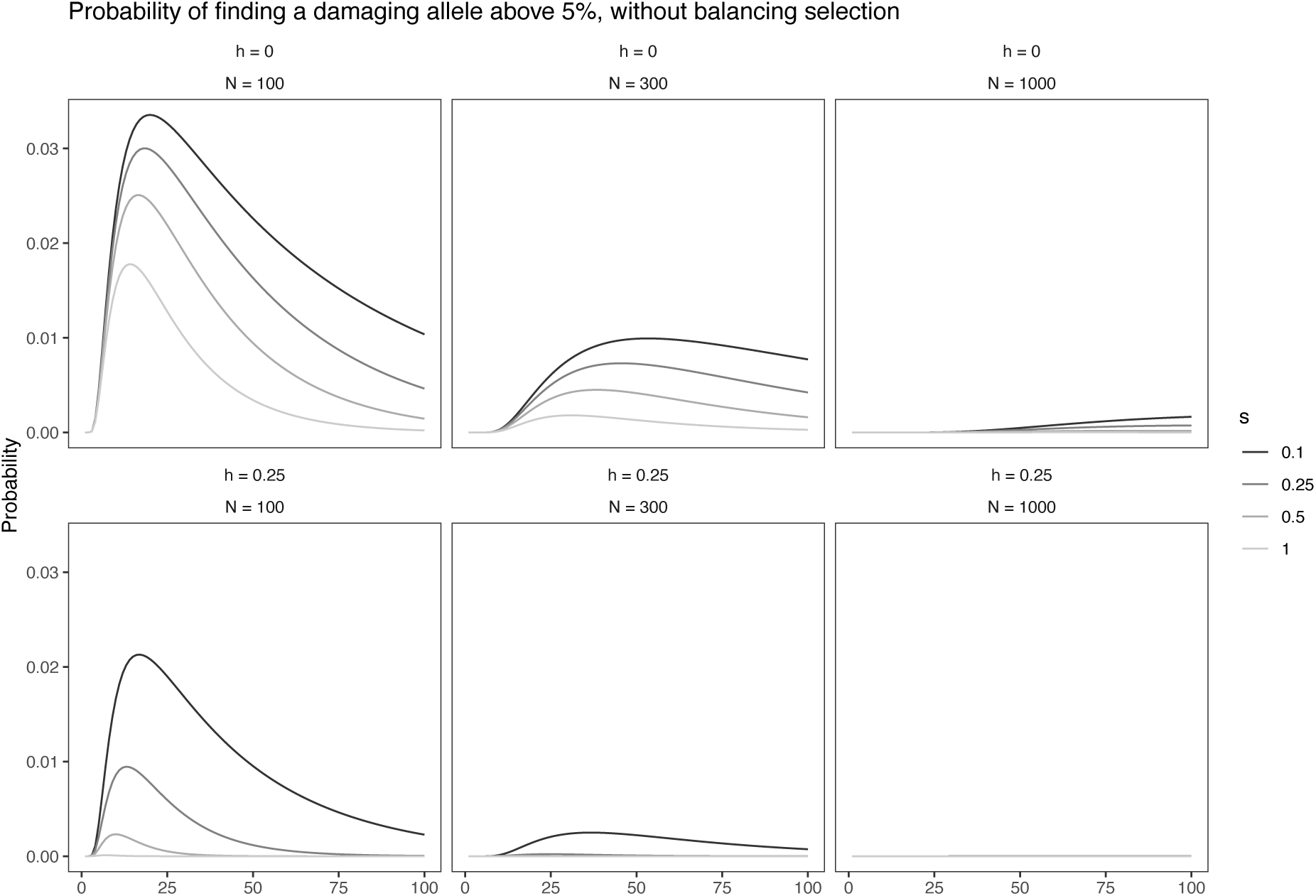
Parameter exploration for damaging alleles without advantage, using the probability matrix model.

**Supplementary Figure 4.**
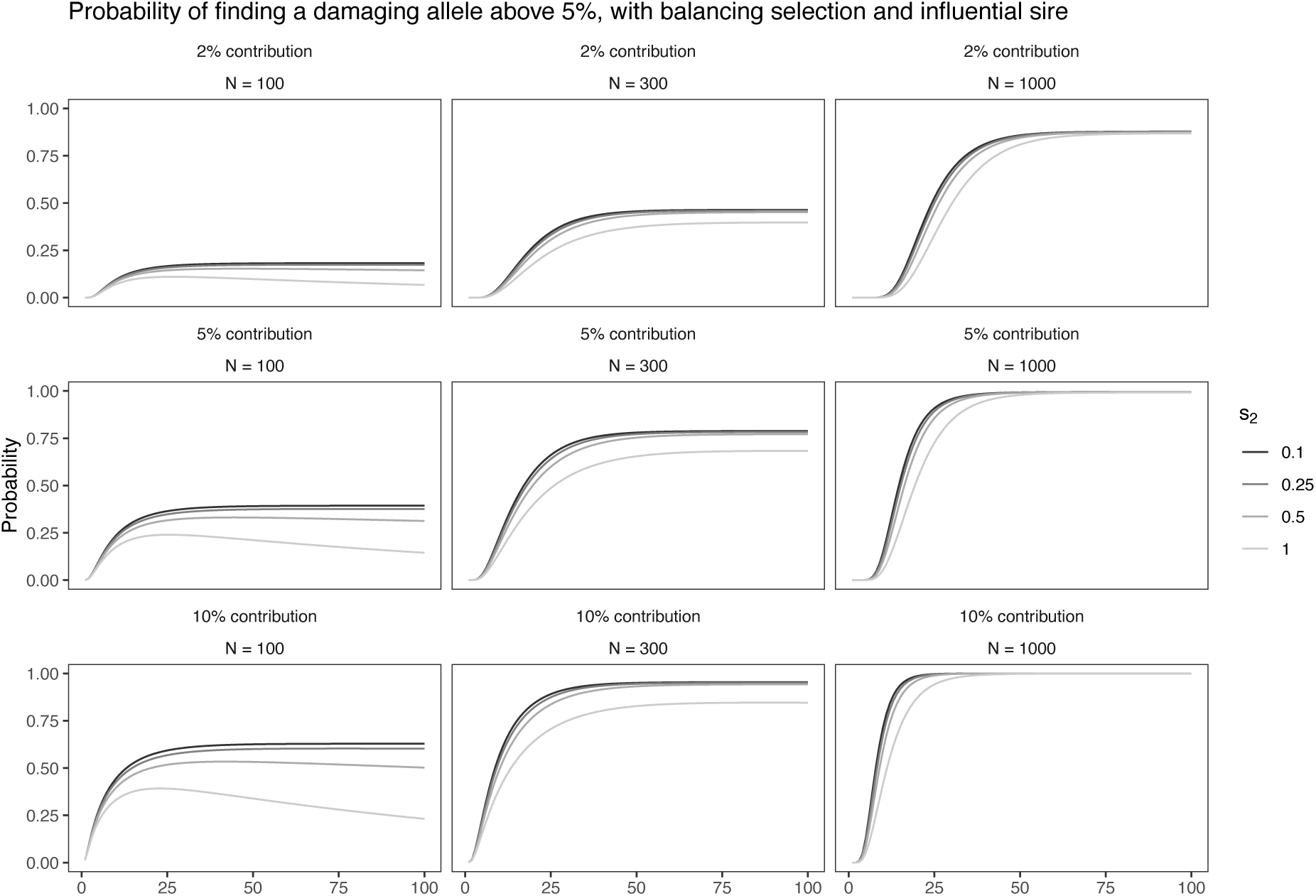
Parameter exploration for damaging alleles with heterozygote advantage after an influential sire event, using the probability matrix model.

**Supplementary Figure 5.**
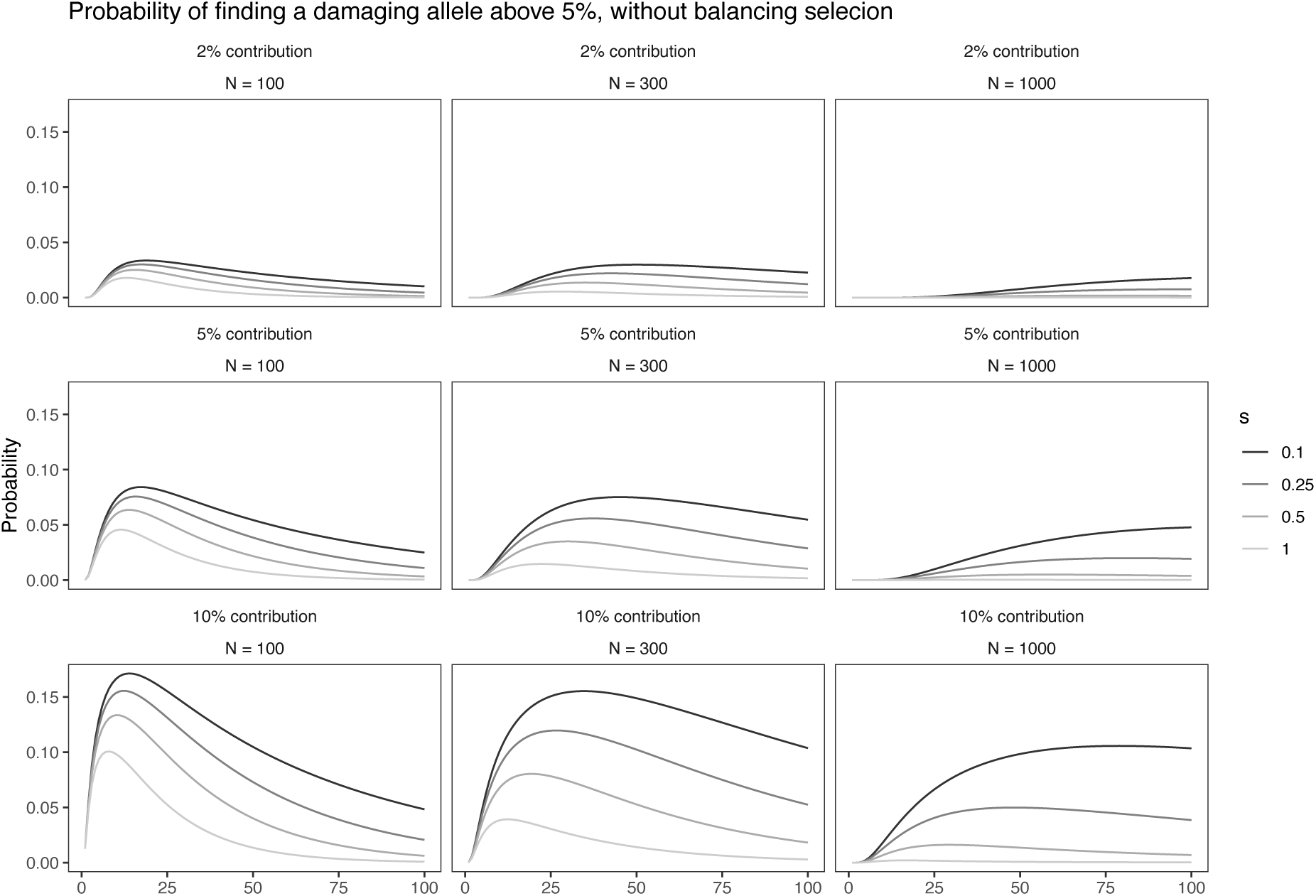
Parameter exploration for damaging alleles without advantage after an influential sire event, using the probability matrix model.

**Supplementary Figure 6.**
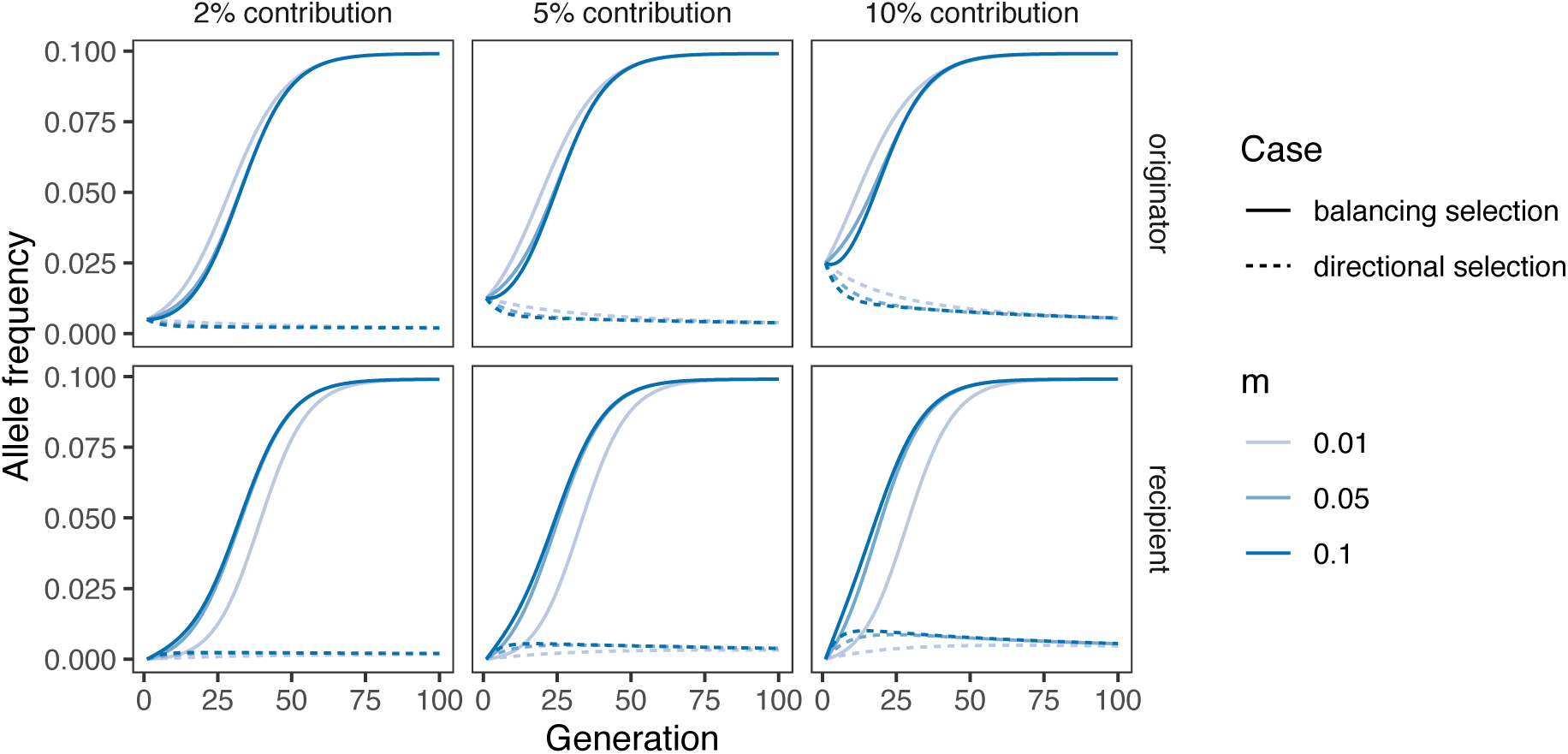
Allele frequency trajectories for lethal alleles with and without balancing selection from the classical deterministic model of a lethal allele with and without heterozygote advantage, in two connected populations with an influential sire event and migration. The colours represent migration rate. Panels show frequency in the originator and recipient populations, and cases with 2%, 5%, and 10% contributions of the influential sire.

**Supplementary Figure 7.**
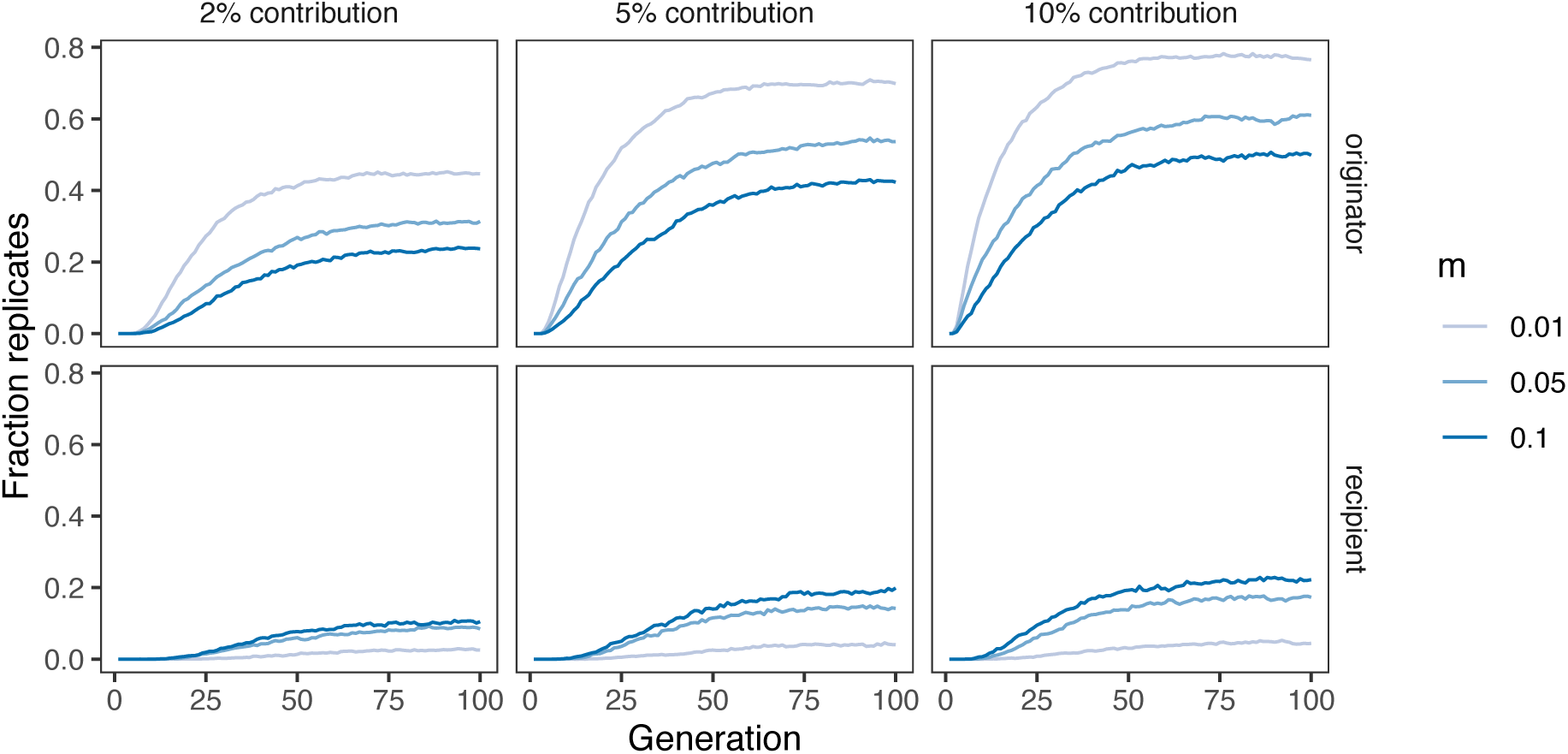
Probability of a recessive lethal allele being observed at high frequency in two populations connected by migration after an influential sire event when there is balancing selection in the originator population but directional selection in the recipient population. The population size of each population was 300. The lines show the fraction of replicates where the lethal allele is at frequency higher than 0.05 in ideal population simulations of two connected populations. The colours represent different migration rates. Panels show frequency in the originator and recipient populations, and cases with 2%, 5%, and 10% contributions of the influential sire.

