## Supplementary Table 1 for "When large-effect damaging alleles enter small populations"

| Case | Reduction time |
| --- | --- |
| Directional selection | 160 |

| Case | Recombination rate | Frequency of beneficial allele in rest of population | Reduction time |
| --- | --- | --- | --- |
| Linked beneficial allele | 0.001 | 0 | 258 |
| Linked beneficial allele | 0.01 | 0 | 246 |
| Linked beneficial allele | 0.1 | 0 | 186 |
| Linked beneficial allele | 0.5 | 0 | 164 |
| Linked beneficial allele | 0.001 | 0.1 | 228 |
| Linked beneficial allele | 0.01 | 0.1 | 220 |
| Linked beneficial allele | 0.1 | 0.1 | 180 |
| Linked beneficial allele | 0.5 | 0.1 | 163 |

| Case | Migration rate | Reduction time |
| --- | --- | --- |
| Directional selection with migration | 0.01 | 111 |
| Directional selection with migration | 0.05 | 115 |
| Directional selection with migration | 0.1 | 118 |
